# Convergent responses to light stress in oligohymenophorean ciliates bearing green algal symbionts

**DOI:** 10.64898/2026.05.24.727488

**Authors:** Joseph Kelly, Niklas Futterknecht, Sven Ernst, Lutz Becks

## Abstract

Photosymbiosis has evolved multiple times independently in ciliates. However, these associations can be antagonized by shifts in environmental parameters that impose stress on the host, necessitating the evolution of mechanisms to contend with this stress and to control the symbiont population. To investigate whether convergent strategies have evolved among algae-bearing ciliates in the class Oligohymenophorea, we imposed light stress on three host species that represent at least two independent evolutionary origins of photosymbiosis and measured their cellular responses. Under high light, all three species experienced an initial drop in host cell density which recovered to levels commensurate with those under low-light conditions as they decreased their symbiont loads. We then performed a comparative transcriptomic study to investigate whether a core set of genes exists that is involved in this response. Thirty-one gene families possess differentially expressed transcripts across all three species that included the upregulation C1 and S28 class peptidases, genes involved in ROS mitigation, and a gene with potential involvement in mitochondrial remodeling associated with changes in algal symbiont load. We additionally found downregulation in *Dicer*, which could mitigate the processing of algal transcripts by the hosts’ RNAi machinery that are freed upon algal digestion, and downregulation of motor proteins that may reflect changes in the host’s swimming behaviors and transport of intracellular vesicles in response to light. The 31 gene families are present and widespread in non-symbiotic oligohymenophoreans, illustrating that a pre-existing genetic toolkit exists in this clade that helps explain how it is predisposed to evolving photosymbioses.

## Introduction

Secondary symbiosis between eukaryotic hosts and photosynthetic microbes has evolved multiple times independently across the tree of life. This phenomenon can have profound effects not only for the host’s physiology as the algae’s photosynthetic byproducts supplement the host’s carbon budget (Erwin & Thacker, 2008; Falkowski et al., 1984), but it can also have effects that ripple throughout the larger community. For example, the secretion of the calcium carbonate skeleton of reef-building corals is enabled by their association with dinoflagellate symbionts (Pearse & Muscatine, 1971). However, given that each participating lineage evolves with regard to its own fitness, antagonizing interactions are inevitable that must be mitigated for the symbioses to be stable. This remains largely an open question in photosymbiotic systems, and our understanding of how this phenotype evolves independently in distantly related clades is in its nascency.

Ciliates (Ciliophora) are among the most intriguing hosts of intracellular photosynthetic symbionts as endosymbiosis with green algae (Chlorophyta) has evolved multiple times independently in this phylum (Esteban et al., 2010). Such ‘green’ ciliates can be found worldwide and constitute a common feature of freshwater planktonic communities. The association is generally understood to be nutritional in nature in that the photosynthates of the algae are used by the otherwise heterotrophic host, such as in *Paramecium bursaria*, which derives maltose from its *Chlorella* and *Micractinium* algal symbionts (Arriola et al., 2018; Brown & Nielsen, 1974). Reflecting the importance of the algae to the host, they are inherited directly by the daughter cells during mitosis, although horizontal transmission has also been demonstrated in the most widely-studied genus of green ciliate, *Paramecium*, when supplied with extracellular algae (Meier & Wiessner, 1988). Ciliates can be specific with regard to their algal symbionts as demonstrated by the host *Tetrahymena utriculariae*, which has only been discovered with the algae *Micractinium tetrahymenae* (Pitsch et al., 2017), and *Paramecium tritobursaria*, which associates with *Chlorella variabilis* (Spanner et al., 2022). However, multiple algal species have also been reported to associate with the same host species (Spanner et al., 2022) and, occasionally, co-inhabit the same host cell (Hoshina & Fujiwara, 2012). Given the pervasiveness of ciliates in Earth’s water bodies and the predisposition of ciliates to establish endosymbiosis with green algae, they constitute an ideal study system to investigate the processes that underlie the independent evolution of this phenotype.

The bulk of what is known about how ciliates control and tolerate their algal symbionts has been determined within the *Paramecium bursaria* species complex. In these species, the perialgal vacuoles housing the symbionts are sequestered at the periphery of the cytoplasm and can be spatially rearranged by the host using a cortical microtubule network (Kodama & Fujishima, 2013; Nishihara et al., 1999). Evidence for positive host control of symbiont populations has been discovered in that the ciliate can allocate nitrogen in the form of amino acids and ammonia to the algae (Albers & Wiessner, 1985; Kato et al., 2006), which promotes their growth while at the same theoretically could provide a means for nutrient embargo.

Negative control of the symbionts is also possible by the host through lysosomal fusion of the perialgal vacuole (Jenkins et al., 2021b; Kodama & Fujishima, 2008, 2013), resulting in the destruction of the algal cell housed therein. However, the algae do appear to have some agency against their demise in that algal transcripts bearing homology to host transcripts are released upon their destruction and are processed by the host’s RNAi machinery, effectively resulting in knockdowns of host transcripts and inhibition of host growth, thus disincentivizing their digestion (Jenkins et al., 2021b). The properties that arise from these mechanisms are consistent with overall stability in the symbiosis, such as the coordination of the division of the algal cells with that of the host cell in which that the algal cell population approximately doubles immediately prior to or in concert with the host cell’s division (Kadono et al., 2004), although perturbations in the environment can shift the direction of interactions between the participating species (Horas et al., 2022).

All organisms must contend with environmental heterogeneity. In photosymbioses, an obviously relevant environmental variable is light intensity, as elevated rates of photosynthesis drive increased production of reactive oxygen species (ROS) (Karpinski et al., 1997). ROS are universally damaging to cells in high concentrations, and hence their overproduction imposes an immediate threat to host fitness. *Paramecium bursaria* appear to have adapted to elevated ROS as evidenced by a higher LD50 of H2O2 relative to *Paramecium* species that do not possess algal symbionts (Kawano et al., 2004). *Paramecium bursaria* also respond behaviorally to elevated ROS and reduce their algal population size (termed symbiont loads) when exposed to high light intensity (Jenkins et al., 2024; Lowe et al., 2016). However, it is yet unknown what genes underlie these adaptations, and whether the same genes are involved across green ciliate species.

In this study we address this question by performing a comparative study of three species belonging to the ciliate class Oligohymenophorea representing at least two independent evolutionary events of the establishment of endosymbiosis with green algae: *T. utriculariae*, *P. tritobursaria*, *and P. deuterobursaria*, with the latter two sister species belonging to the *P. bursaria* species complex (Spanner et al., 2022) that likely inherited the green phenotype from their last common ancestor by the principle of maximum parsimony. Despite being part of the same taxonomic class, the two genera are relatively distant relatives of each other and are separated by 785-1190 million years of evolution, based on fossil-calibrated age estimates of the last common ancestor of *Paramecium* and *Tetrahymena* (Kelly et al., 2025). These host species each possess different algal symbionts, with *T. utriculariae* hosting *M. tetrahymenae*, *P. tritobursaria* hosting *C. variabilis*, and *P. deuterobursaria* hosting *Micractinium conductrix*. We tested the hypothesis that a core set of genes exists among the host species that are involved in coping with light stress and controlling symbiont load by subjecting them to conditions of low and high light intensity. Under high light, all three ciliate species experienced an initial drop in growth rate relative to the low light condition, but upon reducing their endosymbiont load compensated for the detriment in host cell density. A comparative transcriptomic analysis demonstrated that while there is a majority of non-overlapping, differentially expressed orthogroups (gene families) among the three species, there is a core set of orthogroups that are differentially expressed that are candidates for having functional roles involved in coping with stress, such as ROS mitigation and lysosomal activity that could be related to endosymbiont digestion, and which are relevant to changes in the host’s behavior that accompany increased light intensity.

## Results

### Genome assemblies and annotations

*Paramecium tritobursaria* CCAP 1660/21 *and P. duboscqui* PD1 were selected for genome assembly and gene model inference to aid in orthogroup inference and to provide a reference transcriptome for *P. tritobursaria* in the light stress experiment. The assemblies for both species were high-quality based on genome contiguity and the percent of BUSCOs that were recovered as single-copy and complete (Table 1). Likewise, the proteomes inferred from the gene models evidenced that a high proportion of the genes were included in our gene models (Table 1).

**Table 1.** Comparison of *Paramecium* genomes produced in this stud against publicly available genomes. Source information can be found in Table S1.

| Species | Contigs or scaffolds | Total length (Mb) | % GC | N50 (Kb) | No. Proteins | proteome BUSCO |  |  |  |
| --- | --- | --- | --- | --- | --- | --- | --- | --- | --- |
|  |  |  |  |  |  | % Single-copy complete | % Complete duplicated | % Single-copy fragmented | % Missing |
| *P. tritobursaria* | 382 | 33.22 | 29.03 | 110.36 | 17778 | 74.9 | 16.4 | 5.3 | 3.5 |
| *P. duboscqui* | 106 | 27.21 | 31.96 | 378.64 | 11814 | 95.3 | 0.6 | 0.6 | 3.5 |
| *P. deuterobursaria* | 328 | 29.94 | 27.22 | 130.22 | 21220 | 91.2 | 4.1 | 1.2 | 3.5 |
| *P. jenningsi* | 984 | 65.35 | 23.17 | 212.64 | 37098 | 42.7 | 48 | 1.8 | 7.6 |
| *P. tetraurelia* | 697 | 72.10 | 28.04 | 413.03 | 40460 | 29.2 | 69.6 | 0 | 1.2 |
| *P. caudatum* | 1202 | 30.53 | 28.2 | 313.71 | 18673 | 84.8 | 12.3 | 1.2 | 1.8 |

Commensurate with previous knowledge in this phylum, genome-wide phylogenetic inference placed the members of the *P. bursaria* species complex as next closest relatives (Spanner et al., 2022), and supported multiple instances of independent evolution of endosymbiosis with green algae (Figure 1).

**Figure 1.**
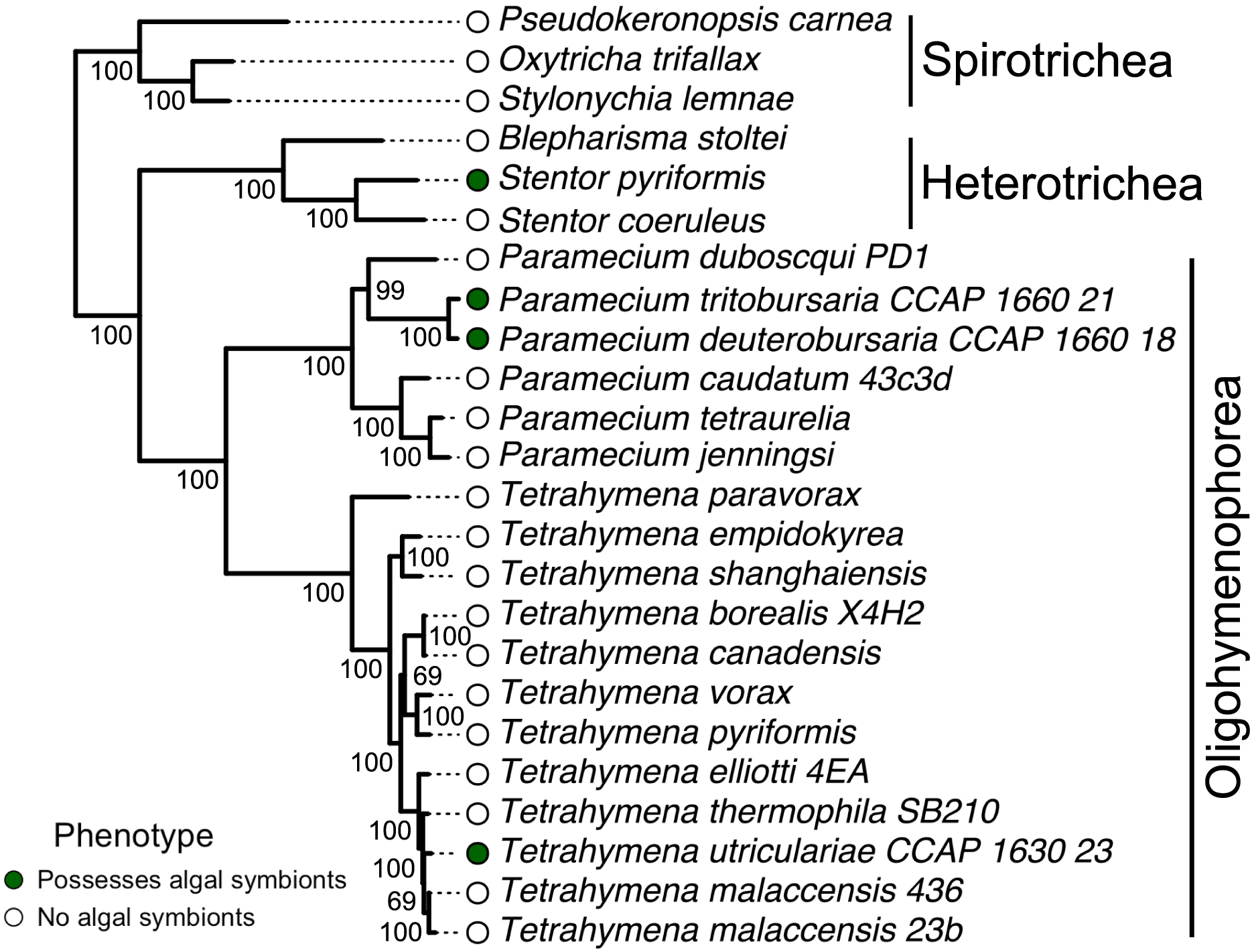
Symbiotic associations with green algae have evolved multiple times independently in ciliates. Phylogenetic reconstruction was performed with IQtree2 using amino acid sequences of 52 alveolate BUSCOs. Bootstrap values are reported as node labels. Clades are annotated to the right of the tips as taxonomic classes.

### Responses of green oligohymenophoreans to light stress

We subjected three species of oligohymenophorean ciliates representing at least two instances of independent evolution of endosymbiosis with green algae to light stress, which is expected to produce elevated levels of ROS due to the algae’s photosynthesis, to investigate whether convergent cellular responses have evolved among these hosts. The three species displayed parallel patterns in culture growth, whereby an initial drop in cell density was experienced in the high light treatment relative to the low light treatment that was compensated for by the end of the experiment (Figure 2, top row). Likewise, all three species exhibited a reduction in chlorophyll autofluorescence under light stress (Figure 2, middle row), which corresponds to a reduction in symbiont load as the mean number of algae per ciliate was found to correlate with Cy5 autofluorescence (linear model: multiple R^2^ = 0.5582, adjusted R^2^ = 0.5072, F-statistic = 10.95, df = 3 and 26, p-value = 7.845e-05). This is corroborated by outright estimates of mean algal cells within each ciliate cell, in which *T. utriculariae* reduced its symbiont load by 30.3% (Welch’s t-test: t = -4.4904, df = 6.4293, p-value = 0.001748), *P. tritobursaria* by 88.2% (t = - 4.0164, df = 4.0442, p-value = 0.007784) and *P. deuterobursaria* by 66.7% (t = -4.8118, df = 6.143, p-value = 0.00139) under high light (Figure 2, bottom row).

**Figure 2.**
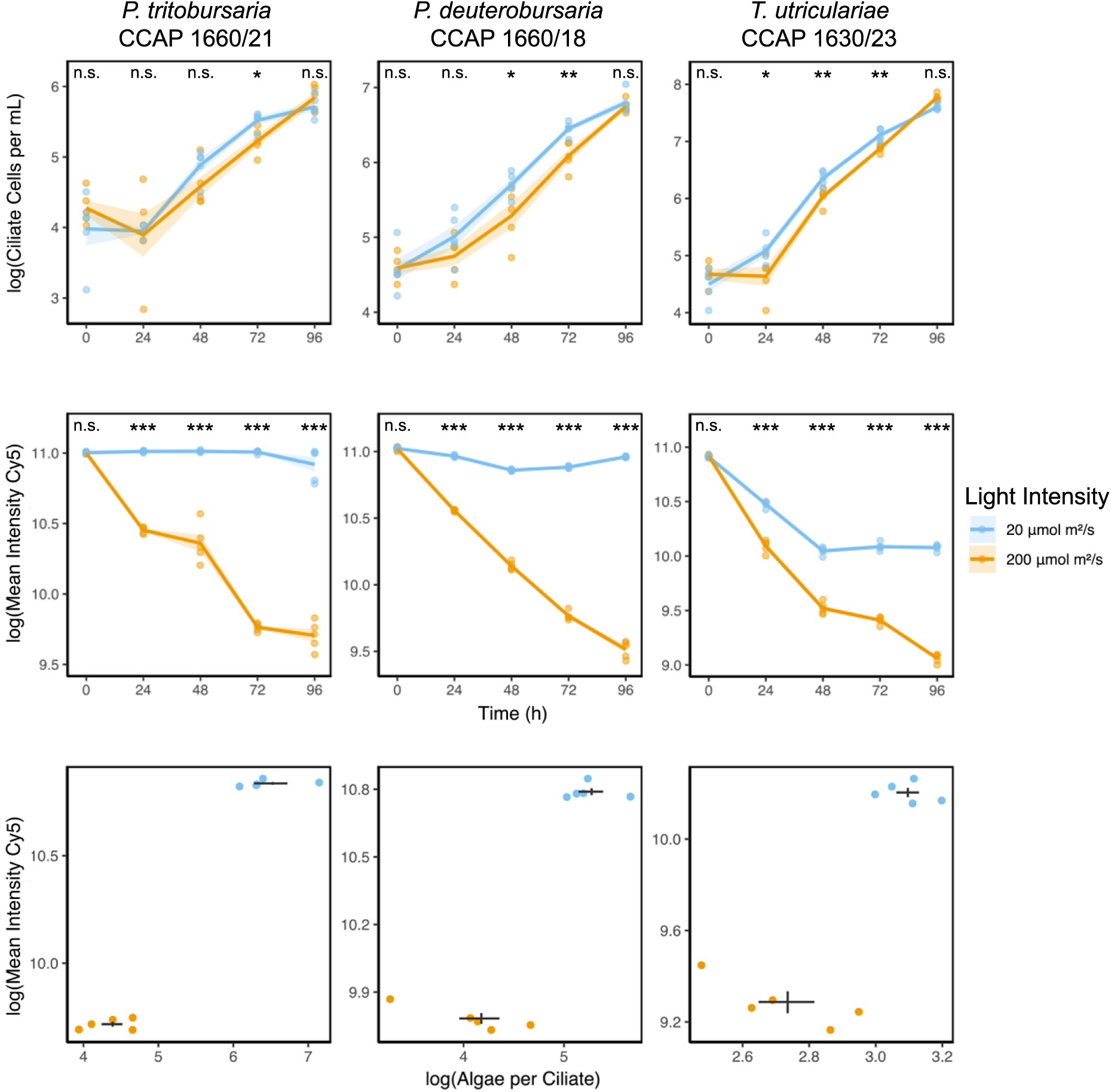
Green oligohymenophorean ciliates exhibit similar phenotypic responses to light stress. Top row and middle rows represent the growth curves of ciliates and the mean intensity of Cy5 channel measuring chlorophyll autofluorescence of endosymbiotic algae within ciliate cells, respectively. The line represents changes of mean values between data points, and the ribbon around this line represents the standard error of the mean. Bottom row: the relationship between chlorophyll autofluorescence values of ciliate cells and the algal symbiont load, measured at the 96-hour timepoint. The sets of intersecting lines depict the standard error of the mean for each measured variable. *** adjusted p-value < 0.001, ** adjusted p-value < 0.01, * adjusted p-value < 0.05, n.s. adjusted p-value > 0.05. N=5 replicates for each species/light treatment combination.

We sequenced the total mRNA fractions of our experimental replicates for all three species at the 96-h timepoint to explore gene expression patterns of the host involved in coping with light stress. The samples segregated in multivariate space by light treatment for each host species, evidencing shifts in transcriptomic profiles (Figure 3A). A greater percentage of genes (24.7%) were differentially expressed in *T. utriculariae* relative to either *P. tritobursaria* (9.6%) or *P. deuterobursaria* (3.6%), and a greater number of orthogroups were found to possess genes that were only differentially expressed in *T. utriculariae* (Figure 3B).

**Figure 3.**
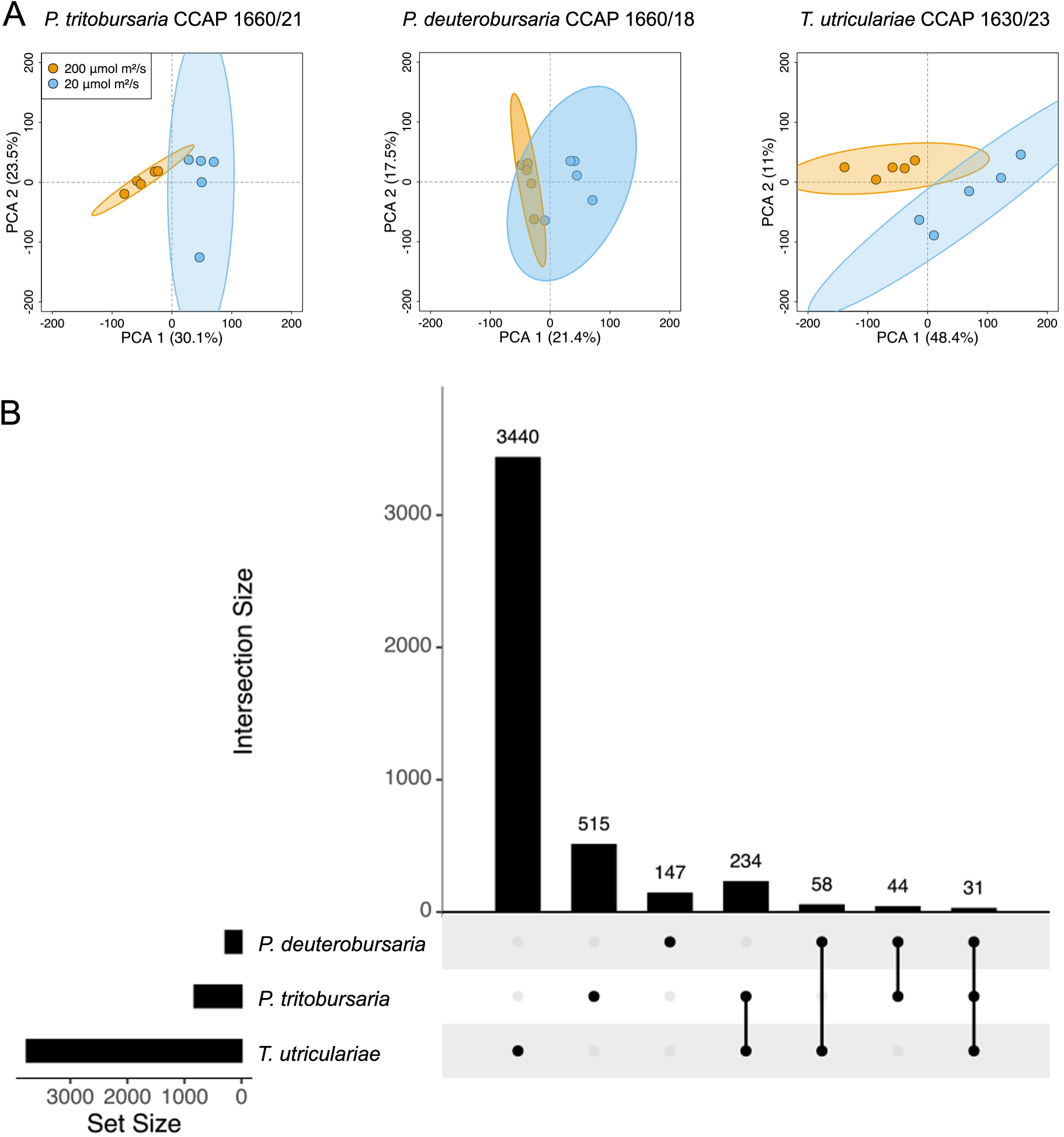
Each of the three ciliate species experiences shifts in transcriptomic profiles during light stress. A: PCAs inferred from the transcriptomes of three ciliate host species. Each point represents a replicate, which are encompassed by 95% confidence intervals **B:** Upset plot showing the intersections among species of orthogroups containing differentially expressed genes, calculated using the entire set of differentially expressed orthogroups for each species.

To examine whether the same orthologs are used by all three species to cope with light stress, we analyzed the overlap in orthogroups that possess differentially expressed members. We found 31 of such orthogroups (Table S1), which was greater than the overlap expected by chance using the Super Exact Test (Wang et al., 2015), a variation of the hypergeometric test for three or more sets (expected overlap = 7.04, fold-enrichment=4.40, p=2.59e-12). This was calculated against the background of the total number of orthogroups that the three host species all had at least one copy in. Twenty-nine of these orthogroups were present in every oligohymenophorean species. OG0001434 (nucleolin, *NCL*, *NSR1*) was missing only in *Tetrahymena borealis* and OG0001518 (alternative oxidase, *AOX*) was missing only in *Paramecium jenningsi*. The differentially expressed members of these shared, ‘core’ orthogroups were found to be enriched for GO terms relevant to motor protein assembly and regulation, and proteolysis (Figure 4). All three host species were enriched for terms GO:0007018 microtubule-based movement, GO:0030286 dynein complex, and GO:0008569 minus-end-directed microtubule motor activity, a pattern driven by differential expression in OG0000012 (dynein axonemal heavy chain, *DNAH*). *Tetrahymena utriculariae* was also enriched for two additional GO terms involved in motor protein activity: GO:0016459 myosin complex and GO:0003774 cytoskeletal motor activity. These GO terms were assigned to OG0000087, which was annotated as dilute class unconventional myosin, a protein family involved in intracellular vesicle transport (MacIver et al., 1998). Both *T. utriculariae* and *P. deuterobursaria* were enriched for GO terms describing proteolytic activity. The first, GO:0008234 cysteine-type peptidase activity was assigned to OG0000003 (Peptidase C1 family) and the second, GO:0006508 proteolysis, was driven by multiple orthogroups including OG0000369 (Serine carboxypeptidase S28 family) and OG0000003.

**Figure 4.**
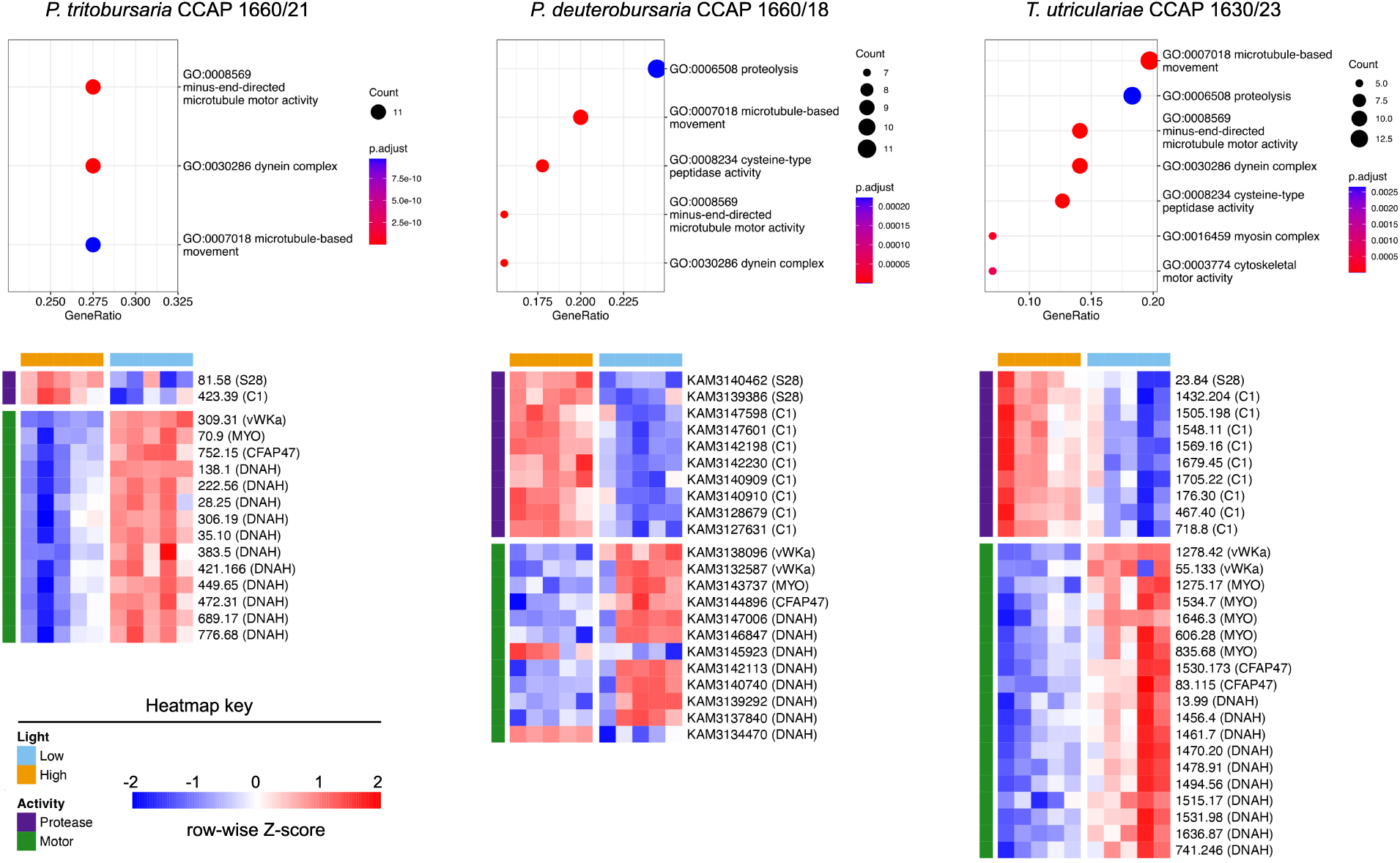
Convergent responses in expression of proteases involved in lysosomal function and motor proteins between green *Paramecium* and *Tetrahymena* to light stress. Top row: Plots of the GO terms that are enriched in the 31 core orthogroups that are differentially expressed across all three host species. Below: Heat maps of key genes involved in driving the enrichment. Each column represents a replicate (n=5 for each species and light treatment combination). Row labels are formatted as ‘transcript.gene’ with the gene symbol in parentheses, with *MYO* abbreviating the dilute class unconventional myosins.

**Figure 5.**
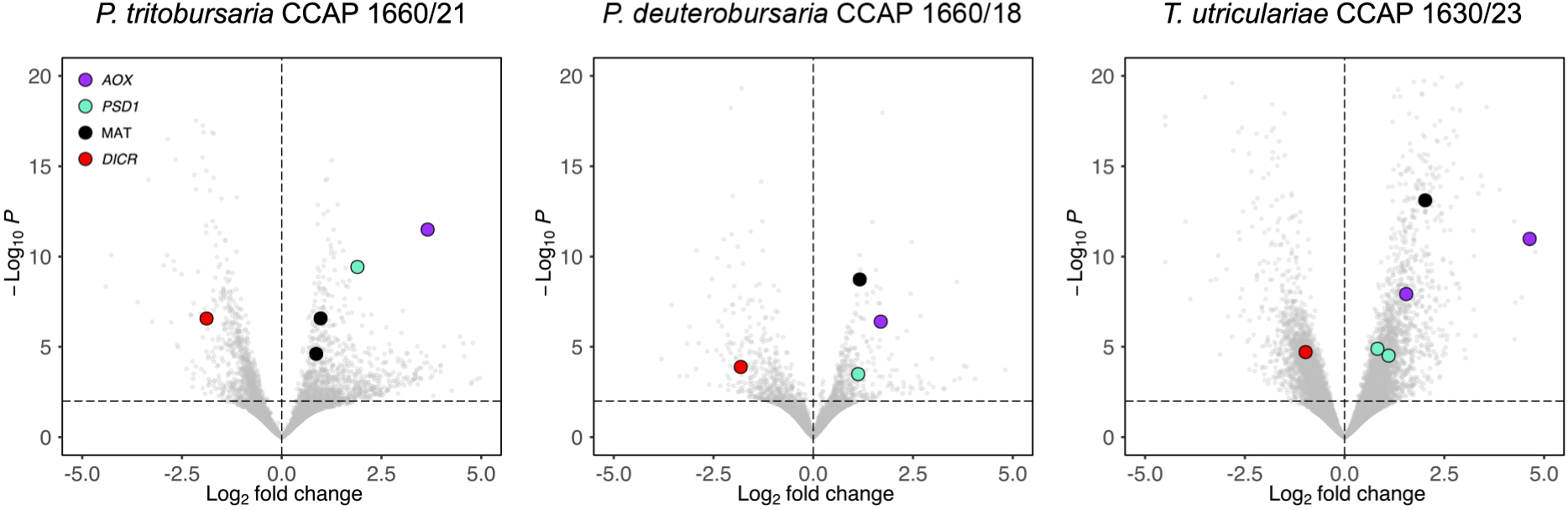
Volcano plots depicting convergent transcriptional responses in genes relevant to ROS mitigation, mitochondrial dynamics, and RNAi among the three host species. Each smaller gray point represents an individual gene. Points above the horizontal line representing the adjusted p-value cutoff of 0.01 of are considered differentially expressed.

We focused our investigation on orthogroups bearing these GO terms and other orthogroups with clear functional annotations that were either collectively or nearly entirely down regulated or upregulated in response to light stress, such that the direction of response was similar across host species and gene copies. Among the orthogroups that fall outside the aforementioned enriched GO terms were alternative oxidase (*AOX*), Dicer (*DCR1*), phosphatidylserine decarboxylase (*PSD*), and S-adenosylmethionine synthetase (*MAT*). *PSD* was further refined to *PSD1*, the paralog which in other species is localized to the mitochondria (Figure S1).

We investigated whether there were any losses or acquisitions of orthogroups that were shared among green oligohymenophoreans. One orthogroup was present only in green oligohymenophoreans and absent in all other species on our tree, OG0025203, which was annotated as *PCSK5* proprotein convertase subtilisin/kexin type 5 [EC:3.4.21.-] in *T. utriculariae* and *P. deuterobursaria*. However, none of the copies were differentially expressed under light stress. They were, instead, expressed at very low levels and were removed during QC steps in the RNAseq pipeline (see Methods). No orthogroups were collectively lost in the three species of interest.

## Discussion

Understanding the genetic components that share involvement across repeated, independent instances of the evolution of a given phenotype can help us identify potential core genes that underly it. We pursued this goal within the context of secondary endosymbiosis between heterotrophic oligohymenophoreans and photoautotrophic chlorophytes with a focus on the hosts given prior evidence of their control over their symbionts (Lowe et al., 2016). We subjected three host species to high light conditions, which would be expected to antagonize the symbiosis given the elevated levels of ROS produced by the algal symbionts. Each species displayed similar phenotypic responses in that they experienced an initial drop in cell density that coincided with a reduction in their symbiont loads by at least a third, although did not experience a net deficit in fitness as evidenced by comparable host cell densities between light treatments at the end of the experiment. This pattern is consistent with an initial stress experienced by the host cells that was dealt with by reducing algal symbiont loads. We examined the transcriptomes of these samples to determine which genes were differentially expressed and whether they belonged to the same gene families (orthogroups) and discovered that a greater number of gene families contained differentially expressed genes than expected by chance. Importantly, each ciliate species we examined hosts a different algal species, and so the cellular responses we observed are not specific to partnership with a single algal strain.

These gene families are present at the root of the Oligohymenophorea, indicating that an ancient, core genetic toolkit exists in this clade that may be responsible for this clades’ predisposition to evolve symbioses with green algae.

### Lysosomes and algal destruction

*Paramecium bursaria* is able to destroy the contents of its parialgal vacuoles by detaching them from the meshwork of cortical microtubules and trafficking them to the lysosome (Kodama & Fujishima, 2008), thus providing a mechanism to control symbiont load. The reduction of intracellular algal populations under light stress has been reported before in *P. bursaria* (Lowe et al., 2016), including in the same strain of *P. deuterobursaria* used in the current study (Jenkins et al., 2024). Thus, our observations of similar responses to reducing symbiont load evidence that a convergent strategy evolved between *P. bursaria* and *T. utriculariae* in dealing with light stress.

We found that one of the most numerous gene families in terms of the number of differentially expressed genes it possesses is the C1 (Papain-like) family of cysteine peptidases, a family of proteases that in humans are primarily localized in lysosomes and other components of the endocytic pathway (Brix et al., 2008) and which are involved in the processing and presentation of antigens derived from an exogenous source, such as pathogens (Villadangos & Ploegh, 2000). In *Drosophila*, the C1 peptidase Cathepsin L (Cp1) is localized in autolysosomes and is required to maintain levels of autophagy during developmentally-triggered midgut degradation and is proposed to be directly involved in digestion of lysosomal cargo (Xu et al., 2021). The role of this gene family has also been described in another protozoan, the kinetoplastid *Leishimania*, whereby inhibition of members of this protein family interfered with autophagy (Williams et al., 2006). Another protease in our dataset, serine carboxypeptidase S28, have similar described properties. In humans, two paralogs are present which are localized to lysosomes (*PRCP*) and other intracellular vesicles (*DPP7*) (Soisson et al., 2010). Orthologs of this gene family are reported to perform digestive functions, such as in trematodes where a lysosomal copy is proposed to be involved in digesting blood meals (Dvorak & Horn, 2018), and in the insect *Tenebrio molitor*, where a member of the gene family is secreted in the midgut of larvae and digests the wheat gluten component gliadin (Goptar et al., 2013). Given that similar activities are reported within each gene family across distantly related organisms, a possible role of these gene families is that that they are involved in digesting the algal symbionts upon fusion of the perialgal vacuole with the lysosome. The differentially expressed genes belonging to these gene families in our ciliate species are collectively upregulated under light stress, consistent with more of these proteins being required during periods of heightened lysosomal activity.

RNAi is a pathway by which cells can protect against unwanted outcomes of mRNA activity and can, for example, act as a mechanism of post-transcriptional gene silencing, protect against viruses, and buffer against transposon replication. The pathway is composed of several conserved proteins and has been demonstrated to be present in both *Paramecium* and *Tetrahymena* (Jenkins et al., 2021b, 2021a; Nekrasova & Potekhin, 2019). In short, the DICR1 endonuclease cleaves algal mRNAs into 23-nt short-interfering RNAs (siRNA) that are then loaded onto the RNA-induced silencing complex (RISC), which uses the siRNAs as templates that anneal to and subsequently cleave target mRNA molecules, hence interrupting their translation. Despite the immunological-like functioning of this pathway, its activity has been demonstrated to be maladaptive to *P. deuterobursaria* in the context of its endosymbiosis with *M. conductrix*. When the *Paramecium* digests the symbiont, algal mRNA molecules that bear homology to the ciliate’s endogenous transcripts are released that are then cleaved by the host DICR1 and processed through the rest of the RNAi pathway, resulting in a knock-down effect of host transcripts. The effect is so potent that the culture’s growth rate can be rescued when symbiont digestion is induced with cycloheximide by knocking down the *DICR1* transcripts themselves (Jenkins et al., 2021b). Under light stress, which induces endosymbiont digestion, we found that *DICR1* is collectively downregulated across the *Paramecium* and *Tetrahymena* hosts, including the same transcript in *P. deuterobursaria* that was targeted in Jenkins et al 2021b. Hence, this phenomenon may be one that is generalizable among these host species and which the ciliate attenuates by downregulating *DICR1*, allowing for a greater capacity to digest algae while still maintaining the host fitness.

### Dealing with ROS

Photosymbiosis is a double-edged association for the host as photosynthesis continuously generates reactive oxygen species (ROS) that can result in nucleic acid damage, protein oxidation, and lipid peroxidation, and are consequently detrimental to cells in excess concentrations (Apel & Hirt, 2004). Importantly, greater light intensities drive increased photosynthetic ROS production, including H2O2 (Karpinski et al., 1997), which is membrane permeable and hence can exit the cell through diffusion, and superoxide, which is exported from cells, including those of chlorophytes (Plummer et al., 2025). Collectively, these processes contribute to the accumulation of ROS in the extracellular environment of phytoplankton (Diaz & Plummer, 2018) which, in the context of intracellular algal symbionts, would translate to ROS accumulation in the cytoplasm of the host ciliate.

Two genes that are upregulated in our host species under high light are relevant to ROS mitigation. The first, alternative oxidase (*AOX*) localizes in the inner mitochondrial membrane and provides an alternative to the cytochrome-based electron transport chain. *AOX* also functions to reduce ROS in mitochondria (Maxwell et al., 1999) and expression of the protein can be induced experimentally by increasing H2O2 concentrations (Wagner, 1995). In both *P. bursaria* (Song et al., 2017) and *T. utriculariae* (Kamal et al., 2025) the perialgal vacuoles are physically associated with host mitochondria, and so the upregulation of *AOX* could be protecting these organelles from H2O2 seeping from the perialgal vacuole. The second upregulated gene relevant to ROS mitigation is S-adenosylmethionine synthetase (*MAT*), which synthesizes S-adenosylmethionine (SAM) that then fed into the transulfuration pathway to produce cysteine, a precursor of the cytosolic antioxidant glutathione (GSH) (Deneke & Fanburg, 1989). While SAM performs multiple roles inside the cell, deficiencies in this compound are linked to elevations in ROS, which are resolved upon supplementation of SAM (Q. Li et al., 2017; Tchantchou et al., 2008). *AOX* and *MAT* upregulation are therefore two genes are candidates for mitigating oxidative stress that could be relevant in different cellular compartments.

### Mitochondrial changes

Multiple studies have reported a relationship between algal symbionts and the ciliates’ mitochondrial shape and number. In *P. bursaria*, mitochondrial densities were reduced in hosts that possessed algal symbionts as opposed to aposymbiotic *P. bursaria* cultures (Kodama & Fujishima, 2022). However, the light intensity used was 20–30 μmol photons m^-2^ s^-1^, approximately that of our low-light treatment and so the dynamics under light stress were not examined. In *T. utriculariae*, aposymbiotic ciliates display rounded mitochondria whereas algal-bearing ciliates displayed dimorphic mitochondria, where those associated with the perialgal vacuole membrane were elongated and the non-associated mitochondria were round (Kamal et al., 2025).

*PSD* converts phosphatidylserine to phosphatidylethanolamine, a phosopholipid that is involved in modifying membrane characteristics. Some eukaryote genomes contain multiple paralogs of *PSD* that possess conserved and distinct functions, where *PSD1* is localized in the mitochondria and contributes to the pool of phosphatidylethanolamine in this organelles’ membrane, whereas *PSD2* is localized in the endomembrane system and contributes to phosphatidylethanolamine in the membranes of other cell components, such as the Golgi apparatus (Voelker, 1997). *PSD1* is required for mitochondrial fusion and its disruption results in defects to the shape of the mitochondria (Joshi et al., 2012; Steenbergen et al., 2005). Given this, it is possible that the upregulation of *PSD1* in host ciliates under light stress is involved in changes to the mitochondria brought on by their interactions with the algae.

### Motor protein expression

Several orthogroups corresponding to motor proteins and their regulatory proteins were nearly entirely downregulated across host species. The first category, relevant to cilia, includes dynein axonemal heavy chain (*DNAH*), the primary force-generating subunit of axonemal dynein (Asai & Koonce, 2001), a motor protein involved in the assembly and function of cilia (Gibbons et al., 2005). Also within this category is the cilia and flagella associated protein 47 (*CFAP47*), mutations of which are implicated in ciliary dysfunction in humans (Ge et al., 2024). *P. bursaria* cells have been observed to decrease swimming activity in lit conditions compared to the dark, whereby the swimming velocity is inversely proportional to light intensity (Matsuoka & Nakaoka, 1988). Given that the primary mode of locomotion of ciliates is cilia, the downregulation of *DNAH* and *CFAP47* likely entail light-dependent, reductive changes in the cilia that reflect changes in their swimming behavior. The second category includes the dilute class of unconventional myosins, which perform intracellular transport in *Drosophila*, mammals, and yeast (MacIver et al., 1998) and a gene involved in the regulation of myosins, *vWFa* (Betapudi et al., 2005; Betapudi & Egelhoff, 2009). Given that some unconventional myosins (e.g. myosin V) transport vesicles along microtubule networks (Conway et al., 2017) it is possible that these genes are involved in the trafficking of the perialgal vacuoles, in which case the cells may be downregulating these gene families simply because there are fewer perialgal vacuoles to organize.

## Conclusion

Investigating the origins of intracellular endosymbiosis and the ways in which it is maintained is central to our understanding of how cellular complexity evolves. To this end, we identified convergent behavioral and transcriptomic responses to light stress in host ciliates bearing algal photosymbionts. We discovered that a core set of gene families are differentially expressed across the host species and are hence candidates for maintaining stability of the symbiosis under light stress. However, light is only one environmental parameter that could destabilize the symbiosis, and our understanding of how the hosts cope with other stressors such as heat, pH, and nutrient limitation could expand the set of gene families that are relevant to host control. Additionally, our transcriptomic survey was taken at only one timepoint, and the nature of the host’s responses is likely different immediately at the onset of the stressor compared to the response after prolonged exposure. Nevertheless, our study elucidated how an ancient set of orthogroups, which are present in the common ancestor of *Tetrahymena* and *Paramecium* and are hence at least 785 million years old, are candidates for being pre-adaptations to symbiosis with green algae and consequently help explain how the ‘green’ phenotype has evolved multiple times in Oligohymenophorea.

## Methods

### Strains used

*Tetrahymena utriculariae* is maintained by the authors and was previously deposited in the Culture Collection of Algae and Protozoa, The Scottish Association for Marine Science, Oban, Scotland, under accession CCAP 1630/23. *Paramecium deuterobursaria* and *P. tritobursaria* were obtained from the same culture collection under accessions CCAP 1660/18 and CCAP 1660/21. *Paramecium duboscqui* PD1 was obtained from the National BioResource Project, Yamaguchi, Japan, under accession, PD000001A.

### DNA isolations, genome assembly and annotation

Cell pellets of *P. duboscqui* PD1 and *P. tritobursaria* CCAP 1660/21 were produced by centrifuging 800 mL of cell culture at 1400 rcf for 5 minutes at 4°C with the rotor break turned off. The resulting cell pellets were quenched in liquid nitrogen and immediately processed with the Wizard HMW DNA Extraction Kit (Promega) following the manufacturer’s instructions for DNA isolation from plant tissue. DNA isolations then underwent HiFi CCS sequencing on the PacBio Sequel II platform following library prep with the SMRTbell Express Template Prep Kit 2.0.

Genome assembly was performed on the HiFi reads using Flye v2.9.1 (Kolmogorov et al., 2019) with two polishing iterations. The contig sets were then processed using PurgeHaplotigs v1.1.2 (Roach et al., 2018) using coverage estimates produced by mapping the HiFi reads to the contigs with minimap v2.24 (H. Li, 2018), an assembly refinement step that has previously been used in *Tetrahymena* to reduce the fraction of micronuclear contamination (Kelly et al. 2025). The final curated assemblies were characterized using Quast v5.0.2 (Mikheenko et al., 2018) and assessed for completeness using BUSCO v5.4.3 (Manni et al., 2021) run on genome mode and specifying the ciliate nuclear code.

Prior to gene annotation, the genome assemblies were soft-masked using RepeatMasker v4.0.7 (Smit & Green, http://www.repeatmasker.org) by combining repeat annotations made by running the program with the *Paramecium* library specified and by performing *de-novo* repeat inference using RepeatModeler v2.0.1 (Flynn et al., 2020). Expressed sequence tag (EST) evidence was produced from cultures of *P. duboscqui* and *P. tritobursaria* that were maintained under 55 μmol/m²/s intensity at constant irradiance. Cell cultures were pelletized as described above and snap-frozen in liquid nitrogen before being stored at -80°C. RNA was isolated and sequenced following Kelly et al., 2025, targeting 100 million read-pairs for each of six *P. tritobursaria* samples and the *P. duboscqui* sample. The raw sequencing reads were then trimmed and filtered to remove low-quality reads using fastp v0.19.6 (Chen, 2023). Algal reads were removed from the *P. tritobursaria* raw reads by mapping them to the *C. variabilis* NC64A genome (NCBI accession GCF_000147415.1) using bbsplit.sh v38.22 (https://sourceforge.net/projects/bbmap/). The read sets for both species were then assembled *de-novo* using Trinity v2.5.1 (Grabherr et al., 2011). Spliced alignments of the transcript assemblies were then made against the corresponding genome assemblies using PASA v2.5.2 (Haas, 2003). A composite evidence approach was then used to infer gene models incorporating expressed sequence tags (EST), publicly available ciliate proteomes, and *ab-initio* gene prediction following the pipeline of Kelly et al., 2025. The resulting proteomes were then assessed using BUSCO run on ‘proteome’ mode and assigned Gene Ontology annotations using EggNogMapper v2.1.12 (Cantalapiedra et al., 2021) and Interproscan v5.55-88.0 (Jones et al., 2014).

### Light stress experiment execution and phenotyping

All experiments and routine cultivation of ciliates were maintained at 20°C under a 14h/10h light-dark cycle using wheat grass medium (doi:10.1101/pdb.rec12079). Prior to experimental set-up, cultures were maintained under 20 μmol/m²/s (the low light treatment) for at least two weeks to allow acclimation. To assemble the experimental replicates, stationary-phase cultures of each of the three ciliate species were passed through cell strainers with 100-micron mesh sizes to remove debris, then subsequently concentrated on cell strainers with 10-micron mesh sizes and rinsed with 30 mL of medium. This step was performed to mitigate autofluorescence from cell debris and clumps of extracellular algae that could otherwise be reported as false positives in the Cy5 channel. Ten replicate cultures of each ciliate species were assembled in 20 mL volumes at 100 cells/mL, then split between two light intensity treatments, a high-light (200 μmol/m²/s) and low-light (20 μmol/m²/s) treatment. Measurements of mean Cy5 intensity and cell densities were obtained on an ImageXpress Micro 4 High-Content Imaging System at 24-hour intervals of aliquots of experimental cultures fixed in 0.1% glutaraldehyde and 0.01% paraformaldehyde (final concentrations). Microscopy images were examined manually to ensure that algae and debris were not counted alongside the ciliates. After the 96-hour intervals were measured, cells were pelletized as reported above and snap frozen in liquid nitrogen before storage at -80°C.

Statistical differences between light treatments were determined within each timepoint using one-tailed Welsh’s t-tests to test the a-priori hypotheses that under high light the host cell densities would be lower and the ciliates would host fewer algal symbionts. The decision to use the Welsh’s t-test was motivated by the observation that many of the comparisons did not meet the assumption of equal variances, as determined using Levine’s test, hence identifying it as preferable to the Student’s t-test. P-values were corrected for multiple comparisons using the Benjamini-Hochberg methodology (Benjamini & Hochberg, 1995).

To examine the relationship between chlorophyll autofluorescence of the ciliates and the number of algae each ciliate cell hosted on average, the experiment was repeated and assayed at the 96-hour timepoint using the IXM to estimate ciliate autofluorescence and a flow cytometer-based assay designed to estimate symbiont load. The latter involved imaging a fixed volume of 100 uL aliquots of experimental cultures on a CytoFLEX (Beckman Coulter) using a gating scheme of R-APC600-A and FSC-A. The gates were established using algal monocultures and cell fractions enriched for ciliates using the cell strainer filtration steps described above.

Both the enriched fractions used to establish the gates and the experimental cultures showed distinct populations with a minimal frequency of intermediate events. Measurements were performed on untreated aliquots and again on aliquots treated with 1% SDS, which bursts the ciliate cells while leaving the algal cells intact. The average number of algae per ciliate was estimated as the ((number of algae in the SDS-treated samples) – (number of algae in non-SDS treated samples))/(number of ciliates in non-SDS treated samples). Flasks were re-randomized daily within each light level for each experiment. We used a linear model to test whether the average number of algae per ciliates was correlated to the mean intensity of the Cy5 channel of the ImageXpress Micro 4 High-Content Imaging System using a linear model with species as a fixed effect using the following syntax: lm(Cy5_intensity ∼ algae_per_ciliate + species).

### Transcriptomic analysis of light stress experiment

Cell pellets of the light stress experiment were resuspended in 1 mL of Trizol, transferred to prefilled 2mL tubes containing 0.1 mm zirconium beads (Merck, cat. Z763764), and homogenized at 4200 rpm for 1 minute on the PowerLyzer 24 Homogenizer (Qiagen). Phase separation was performed following the manufacturer’s instructions. The aqueous phase was then mixed in a 1:1 ratio with 100% isopropanol and transferred to the columns of the RNEasy mini kit (Qiagen). RNA was then cleaned following the manufacturer’s instructions, including the DNase I on-column digest step. Sequencing libraries were then produced following poly-A enrichment and sequenced using a 2×150 paired-end chemistry on an Illumina NovaSeq6000, targeting 30 million read-pairs per sample.

Sequencing reads were then processed with fastp as described above. Transcript abundance was then estimated using salmon v1.10.3 (Patro et al., 2017) run on mapping-based mode. The resulting quant.sf files were then imported into R v4.2.0 (R Core Team, 2022) using the ‘tximport’ function of the tximport v1.24.0 package (Soneson et al., 2015), and converted to a DESeqDataSet object using the ‘DESeqDataSetFromTximport’ function of DESeq2 v1.36.0 (Love et al., 2014). Low-expression genes (less than ten transcripts across six or more samples) were removed prior to downstream analysis. Log-fold changes in expression values between the low and high light treatments were calculated using the ‘lfcShrink’ function of DESeq2 with the ‘apeglm’ shrinkage estimator. PCAs were constructed by normalizing the counts in the DESeqDataSet object using the ‘counts’ function specifying ‘normalized=T’, then by processing the matrix with the ‘prcomp’ function in the ‘stats’ package. Orthogroups that contained differentially expressed genes, defined as possessing an adjusted p-value < 0.01, were identified and overlap of differentially expressed homologous transcripts were visualized using the ‘upset’ function of the ‘UpSetR’ package (Conway et al., 2017). Genes were then identified that belonged to orthogroups that are differentially expressed and belong to the 31 orthogroups that contain differentially expressed genes in all three species. The enrichment of GO terms in these sets of genes was inferred using the EggNogMapper and Interproscan annotations described above and the ‘enricher’ function of the R package clusterProfiler v4.4.4 (Wu et al., 2021) using a p-value cutoff of 0.01 and a minimum gene set size of 20.

### Phylogenetic analysis and orthogroup inference

Publicly available proteomes of *Paramecium* and *Tetrahymena* species as well as proteomes belonging to the ciliate classes Spirotrichea and Heterotrichea were compiled for the sake of phylogenetic reconstruction and orthogroup inference (Table S2). BUSCO was performed on the proteomes using the alveolata_odb10 database. A set of 80 initial BUSCOs present in at least 80% of species were then used for phylogenetic inference. Alignments of peptide sequences within each BUSCO were made with Muscle v5.1 (Edgar, 2004) and subsequently filtered using Gblocks v0.91b (Castresana, 2000) with ‘-b5=h’ specified. Model selection was then performed on alignments with at least 50 positions using ModelFinder (Kalyaanamoorthy et al., 2017) implemented in IQtree v2.2.3 (Minh et al., 2020). Any alignments failing the composition chi^2^ test were omitted from further analysis, resulting in a final dataset of 52 BUSCOs. Phylogenetic inference was then performed using IQtree2 using an analysis partitioned across the BUSCOs and by performing 10,000 ultrafast bootstrap replicates (Hoang et al., 2018). Orthogroups were then inferred among the full set of taxa for the sake of identifying homologs among the three focal species by supplying the proteomes and phylogeny, which was rooted on the Spirotrichea, to orthofinder v2.5.4 (Emms & Kelly, 2019).

An additional phylogenetic analysis was performed on the differentially expressed members of OG0001367, which was annotated as *PSD*. This was motivated by the fact that multiple PSD paralogs were recovered for each species. *PSD* is a gene family where the different paralogs have distinct and conserved functions, and so we sought to refine our functional annotations to identify what the potential roles of the ciliate copies might be. Gene tree inference was performed as described above by aligning the sequences with MUSCLE, filtering the alignment with Gblocks, and by running IQtree2 with 1000 ultrafast bootstrap replicates.

## Data Availability

Raw sequencing data for this study are available under NCBI BioProject ID PRJNA1465301. Phenotyping datasets and genome assemblies and annotations are available at Zenodo doi: 10.5281/zenodo.20358100.

## Acknowledgements

This material is based upon work supported by the NSF Postdoctoral Research Fellowships in Biology Program under Grant No. 2109477 awarded to JBK, a grant from the Young Scholars Fund of the University of Konstanz awarded to JBK, and by a grant made by the Gordon and Betty Moore Foundation to LB (grant no. 9196; https://doi.org/10.37807/GBMF9196). We acknowledge the flow cytometry center (FlowKon) at the University of Konstanz for the use of flow cytometers and the expert support in instrument usage and data analysis. The authors would like to thank the Stony Brook Research Computing and Cyberinfrastructure and the Institute for Advanced Computational Science at Stony Brook University for access to the high-performance SeaWulf computing system, which was made possible by $1.85M in grants from the National Science Foundation (awards 1531492 and 2215987) and matching funds from the Empire State Development’s Division of Science, Technology and Innovation (NYSTAR) program (contract C210148).

## Supplementary Figures

**Figure S1.**
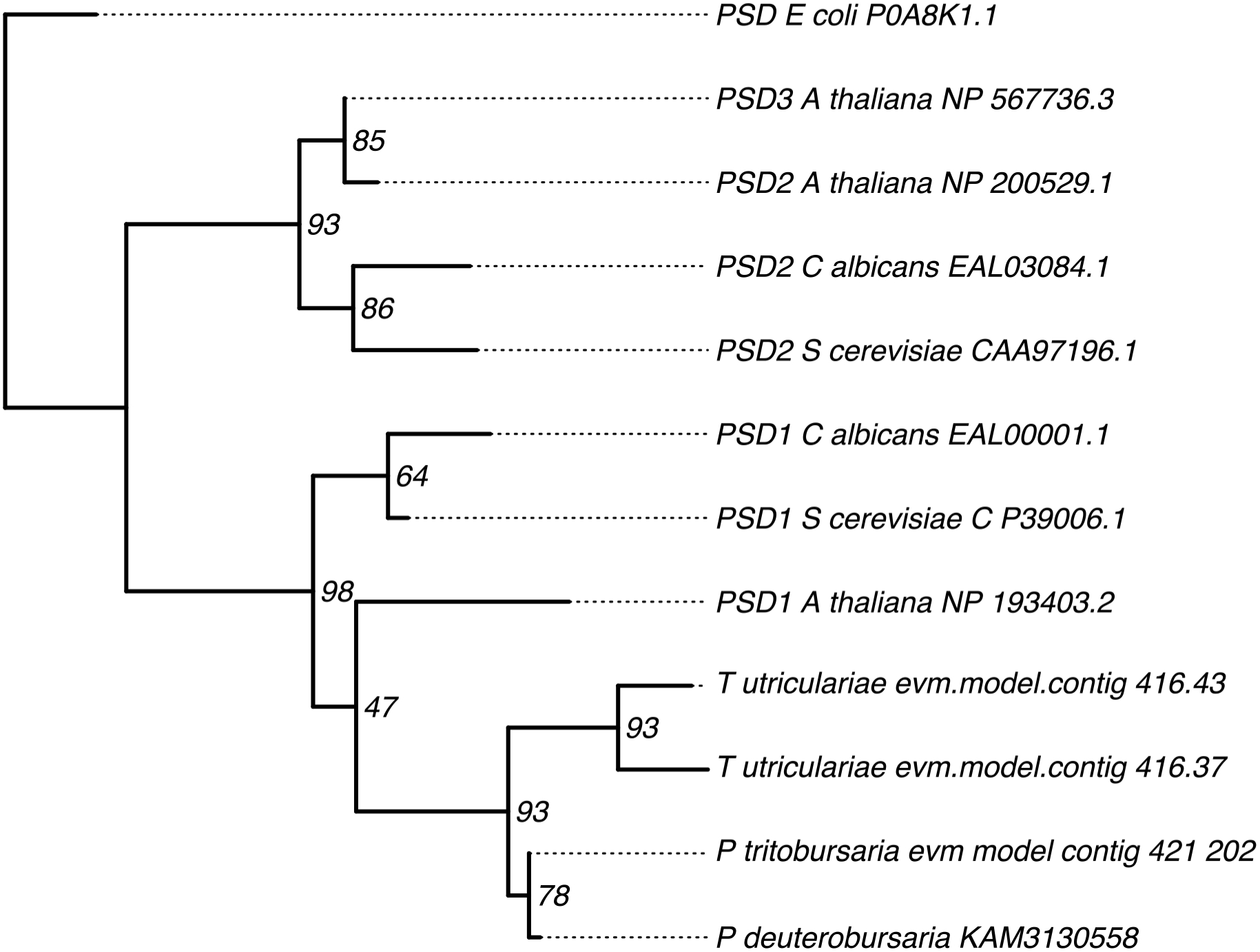
Gene tree of publicly-available *PSD* homologs and the differentially expressed members of OG0001367, rooted on the *E. coli* copy. Node support values are bootstrap support values.

## Notes

### Competing Interest Statement

The authors have declared no competing interest.

